# LMM-MQM time series mapping - An application in a murine advanced intercross line identifies novel growth QTLs

**DOI:** 10.1101/2022.01.23.477441

**Authors:** Danny Arends, Deike Hesse, Stefan Kärst, Sebastian Heise, Shijie Lyu, Paula Korkuc, Manuel Delpero, Megan K. Mulligan, Pjotr Prins, Gudrun A. Brockmann

## Abstract

The Berlin Fat Mouse Inbred line 860 (BFMI860) is a mouse model for juvenile obesity. Previously, a recessive major effect locus (*jObes1*) was identified on chromosome 3 explaining around 26% of the body weight variance in an BFMI860xC57BL/6NCrl advanced intercross line. The aim of this study was to discover additional QTL.

Time series body weight data were modeled using linear mixed models (LMM), while a multiple QTL mapping (MQM) approach compensated for the *jObes1* locus effect. LMM-MQM identified five additional loci significantly associated with body weight. Variance explained by the *jObes1* locus increased to 38.1% when using LMM-MQM mapping, while the additional loci explained between 2.0% and 3.9% of the body weight variance. Several positional candidate genes within the novel QTL regions were found in KEGG pathways for insulin signaling and insulin resistance. Strong distortion with preference for the BFMI allele was observed within a newly identified QTL containing the well-known *Foxo1* regulator of adipocyte differentiation.

Here, we present a novel method for QTL detection in time series data: LMM-MQM time series mapping. We show that our method is more powerful in detecting QTLs compared to single timepoint mapping approaches. Thus, the time series structure should be considered for optimal detection of small effect QTLs. LMM-MQM time series mapping can be used to find genetic determinants of all kind of “phenotypes over time” be it lactation curves in cattle, plant biomass, drug clearance in human clinical trials, or cognitive decline during disease.

## Introduction

Weekly body weights are often collected during animal quantitative trait locus (QTL) experiments to monitor animal health and as an additional phenotype for mapping. When confronted with time series data, QTL mapping is normally performed at each time point independently ^1–3^. However, QTL mapping per time point fails to benefit from the underlying repeated measurement structure of time series data ^4,5^. Additionally, mapping time points independently does not take into account that genetic effects can be time-dependent with effect sizes varying over time ^6^, this can lead to failure to detect loci of interest or underestimation of variance explained by a given locus.

Recently, more sensitive statistical methods have been developed for QTL mapping, such as composite interval mapping (CIM) ^7^, multiple QTL mapping (MQM) ^8,9^, LASSO ^10^, Bayesian mapping methods ^11^, and machine learning ^12^. These methods aim to provide a better dissection of genetic effects compared to classical interval mapping ^13^ by using more advanced statistical modeling, prior information and/or incorporating genetic cofactors to compensate for known genetic effects. While, these methods are generally simple and computationally efficient they are limited to mapping individual time points.

Statistical methods and approaches have also been developed to model time series data. The simplest approach is to take the average or maximum LOD score ^14^ across the time series. More complex methods rely on linear mixed models (LMM) ^15,16^ to account for repeated measurements and covariance structure, or employ a two-step modeling approach ^17^ estimating growth curves for each individual and then associating curve parameters with genetic markers ^18^. While these methods allow to model temporal data, they are not designed to handle genetic loci known to affect the phenotype under investigation.

The Berlin Fat Mouse Inbred line (BFMI), a mouse model for juvenile obesity ^19,20^, which becomes obese under a standard maintenance diet at around 42 days ^20^ and acquire several features of the metabolic syndrome including reduced insulin sensitivity ^21^ at around 70 days. Previously, individual time point mapping of growth data from BFMI identified a major effect locus on chromosome 3 (*jObes1*) ^22^, which besides body weight also pleiotropically affects the ratio between fat and lean mass. An advanced intercross line (AIL) between the obese BFMI860 and the lean reference mouse strain C57BL/6NCrl (B6N) showed that the *jObes1* locus explains 25.6% of the variance in body weight. The *jObes1* locus combined with other sources of variance: subfamily (25%), litter number (6%), litter size (5%), and seasonal effects (1%) still leaves around 37 % of body weight variance unexplained ^23^. Therefore, more loci are expected to significantly contribute to the body weight of the AIL population.

Here, we present a novel method called LMM-MQM time series analysis that combines the strengths of two well established (QTL mapping) methods to discover additional genetic loci associated with body weight across different time points. We use MQM to compensate for known genetic loci ^8,9^ and combine this with the flexibility of modeling time series data using LMMs to consider random effects and repeated measurements ^24^. The method presented here is generic and can be used to perform association analysis of any phenotype measured over time when genotypes are available.

## Material & Methods

### Mouse population

This study is based on genotypic and phenotypic data of 344 male mice of an advanced intercross line (AIL) in generation 28. The AIL population was generated from the mapping population of a cross between a male mouse of the obese line BFMI860 and females of the lean reference mouse strain C57BL/6NCrl (B6N). Beginning in generation F_1_, individuals from the same generation were randomly mated using the program RandoMate ^45^.

### Husbandry conditions

Mice were maintained under conventional conditions and a 12:12 h light:dark cycle (lights on at 6:00 a.m.) at a temperature of 22 ± 2°C. Animals had *ad libitum* access to food and water. They were fed with a rodent high-fat diet (HFD) containing 19.5 MJ/kg of metabolizable energy, 45% from fat, 24% from protein and 31% from carbohydrates (E15103-34, ssniff EF R/M, Ssniff Spezialdiäten GmbH, Soest / Germany). All experimental treatments of animals were approved by the German Animal Welfare Authorities (approval no. G0065/14).

### Phenotypes

Body weight of AIL animals was recorded weekly between the age of 21 and 70 days. Phenotypes and covariates can be found in supplemental file 1.

### Genotypes and SNP quality control

Genotypes of 344 males of AIL generation 28 were generated at GeneSeek (Lincoln, NE) using the second-generation Mouse Universal Genotyping Array (MegaMuga, Illumina, San Diego, CA). This array contains probes targeting around 77,800 known SNPs. Probes were remapped to the mouse reference genome (GRCm38) using BLASTN ^46^. Probes that did not map uniquely were removed from further analysis. SNP genotype calls with low confidence (< 0.9) were set to missing. Furthermore, when a SNP had less than 30 individuals in a genotype group, that genotype group was set to missing to prevent false association. After this procedure non-segregating SNPs were removed. Furthermore, SNPs were discarded when 1) the founder information was not available 2) one (or both) of the founder strains were heterozygous or 3) founders have the same genotypes. In total 17,971 SNPs passed these very stringent quality checks. Genotype data are available in supplemental file 1.

### Modeling of time series data

The individual’s body weight as phenotype response of all models is coded as P. In the case of per time point linear model (LM) and linear mixed model (LMM) mapping this is the body weight at the time point under consideration. In LMM-MQM time series mapping P stands for all body weight measurements obtained during the entire experiment. Fixed and random effects are encoded using the following abbreviations:

- F = Family structure, represented by the father of the litter (31 groups)
- L_s_= Litter size, the number of animals born in a single litter (5 groups: 8, 9, 10, 11, and 12 offspring)
- L_n_= Litter number, the n^th^ litter of a mother (5 groups: A, B, C, D, and E). Considering biological evidence, a two-group encoding was also tested during the model phase to differentiate the first litter of a mouse from the following litters phase (2 groups)
- L_t_ = Litter type, the combination of litter number and litter size. Depending on the coding of the litter number this leads to a different number of factor classes for the litter type. L_t2_ means litter type encoded using litter number as two levels (resulting in 8 groups), L_t5_ stands for litter number encoded using the original five levels (resulting in 14 groups)
- S = Season of birth (3 groups: Fall, Spring, and Winter)
- T, T^2^ and T^3^ = Time of measurement (8 timepoints day 21, 28, 35, 42, 49, 56, 63, and 70 modeled as a linear effect shifted by 21 days, making the day 21 the intercept)
- M_(*jObes1*)_ = Genotypes at the top marker of the *jObes1* locus (3 groups: B6N/B6N, B6N/BFMI, and BFMI/BFMI)
- M_(x)_ = Genotypes at the marker under consideration, this can be 2 or 3 groups depending on the number of unique genotypes (AA, H, BB) at a certain marker
- R_(1|S)_ = Random subfamily effect: Subfamily as grouping factor, no slope parameter
- R_(1|Ind)_ = Random individual effect: Individual as grouping factor, no slope parameter
- R_(Time|Ind)_ = Random effect: Individual as grouping factor, time as slope parameter
- E = Error term of the model

Before modeling, it was essential to reduce the complexity of the model and to find an optimal strategy to deal with cofactors that contain a large number of groups, in our case: litter number and litter size. From experience we know that there is a significant difference in litter size and litter weight between the first litter of a female compared to following litters by the same female. We defined L_n2_ and L_n5_, encoding for litter number as two (1^st^ litter versus next litters) or five levels (litter A to E), respectively. Model selection (supplemental file 2) showed that including litter number and litter size was done most parsimonious when coded as litter type with two levels for litter number (L_t2_) which results in 8 groups.

Body weight was modeled by including the largest environmental covariate (subfamily) found during our previous work, creating an initial null-model for comparison. Afterwards, all other environmental covariates were included in a stepwise fashion, adding covariates to the model if they significantly improve upon the previous model (>10 AIC units drop) (Table 1). When no environmental covariates were left to include or exclude from the model, time was tested as a fixed effect. After including time as fixed effect it was tested as the slope parameter in the random effect model. Modeling was continued by adding higher exponents of time into the model as fixed effects, adding additional inflection points to the model to improve the fit of the model ^4^, using the 10 AIC unit drop criterion for inclusion into the model.

**Table 1:**
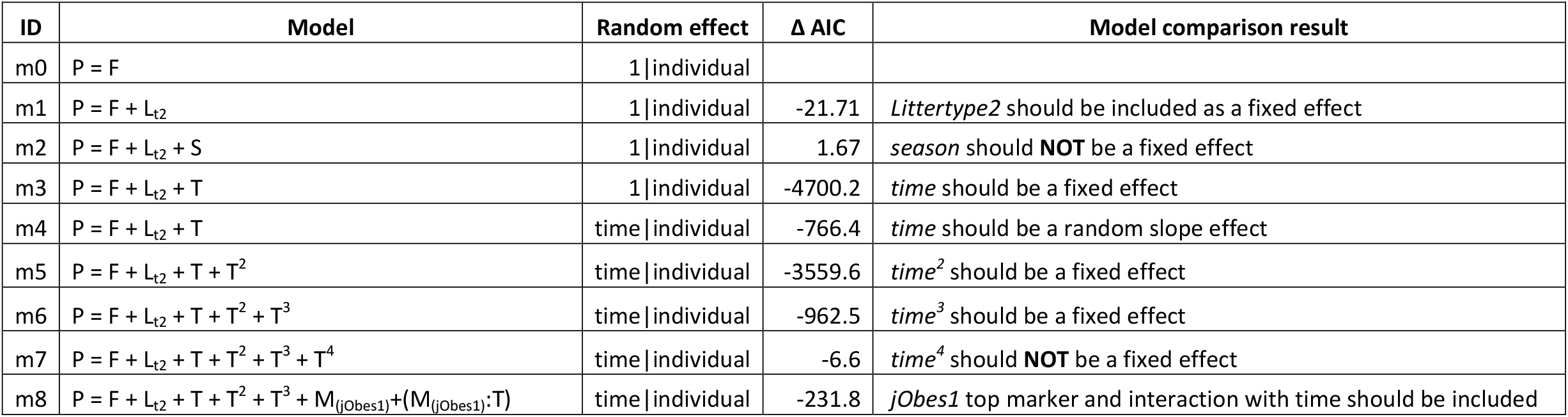
Time series modeling steps to obtain a null-model to use for LMM-MQM mapping. Starting with a model including Family structure and litter type environmental covariates are added to the model and their Δ AIC is calculated relative to the previous best model. Including season actually decreases the fit of the model relative to the amount of degrees of freedom lost. The conclusion/result from each modeling step is summarized in the last column. Modeling the available time series data resulted in m10 being the most suitable model, and as such it is used in the LMM-MQM mapping phase as the null-model.

| ID | Model | Random effect | $\Delta$ AIC | Model comparison result |
| --- | --- | --- | --- | --- |
| m0 | $P = F$ | 1 \| individual | | |
| m1 | $P = F + L_{t2}$ | 1 \| individual | -21.71 | *Littertype2* should be included as a fixed effect |
| m2 | $P = F + L_{t2} + S$ | 1 \| individual | 1.67 | *season* should **NOT** be a fixed effect |
| m3 | $P = F + L_{t2} + T$ | 1 \| individual | -4700.2 | *time* should be a fixed effect |
| m4 | $P = F + L_{t2} + T$ | time \| individual | -766.4 | *time* should be a random slope effect |
| m5 | $P = F + L_{t2} + T + T^2$ | time \| individual | -3559.6 | *time*<sup>2</sup> should be a fixed effect |
| m6 | $P = F + L_{t2} + T + T^2 + T^3$ | time \| individual | -962.5 | *time*<sup>3</sup> should be a fixed effect |
| m7 | $P = F + L_{t2} + T + T^2 + T^3 + T^4$ | time \| individual | -6.6 | *time*<sup>4</sup> should **NOT** be a fixed effect |
| m8 | $P = F + L_{t2} + T + T^2 + T^3 + M_{(jObes1)} + (M_{(jObes1)} \cdot T)$ | time \| individual | -231.8 | *jObes1* top marker and interaction with time should be included |

When no more exponentiations of the time parameter are left that significantly improved the model, genetic loci known to influence the AIL body weight were included. Genetic covariates in the BFMI AIL population consist of the *jObes1* locus on chromosome 3 (represented by the genotype at top marker) that had been detected before. This locus was added into the model and tested for its ability to improve the fit of the model. The resulting model was defined as the null-model for LMM-MQM time series mapping.

### QTL mapping

Standard QTL mapping for each of the 8 measured time points was performed using a standard linear model (LM) (Results published before ^23^) and a standard linear mixed model (LMM). The M_(jObes1)_ term was only included in the model when performing multiple QTL mapping (MQM) ^9^ to correct for the *jObes1*. Additionally, this the M_(jObes1)_ term was dropped when associating markers close (+/- 5 Mb) to the *Jobes1* locus.

The following linear model (LM) that was used per time point includes F as family structure which is represented by the father of a litter to absorb population structure:

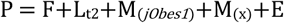

The following linear mixed model (LMM) used per time point additionally includes subfamily as a random effect to absorb any residual population structure, not absorbed by F:

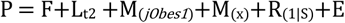

### LMM-MQM time series mapping

Modeling of time series resulted in the following linear mixed model that was fitted to the growth phenotype across all eight time points (For modeling steps see Table 1). Time points were shifted by 21 days, such that the body weight measurements correspond to “days after first weighing”. The model includes the previously identified *jObes1* locus as a main effect covariate M_(jObes1)_ and as a covariate-time interaction M_(jObes1)_:T. Since the *jObes1* locus could directly affect the body weight (M_(*jObes1*)_) as well as cause differences in body weight gain over time (M_(*jObes1*)_:T) throughout the experiment. Derivation of the null-model by AIC selection (Table 1), leads to the following null-model for mapping QTL for the growth curves:

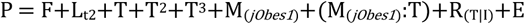

LMM multiple QTL mapping assesses the direct effect of the marker M_(x)_ under investigation on the body weight, but again is also allowed to affect body weight gain over time. This is done by including the interaction term (M_(x)_:T) into the model, leading to the following full-model including the marker effect:

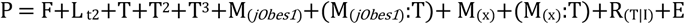

The full-model at each marker is tested against the null-model using the built in ANOVA χ^2^-test available from the R language for statistical computing ^48^. Resulting p-values were converted to LOD scores by taking the −log_10_(p-value). When mapping within 5 Mb of the *jObes1* top marker on chromosome 3, the M_(*jObes1*)_ and (M_(*jObes1*)_:T) terms are dropped from the model as is required in multiple QTL mapping ^8,9^.

Since many markers are in strong linkage disequilibrium (LD) with each other in AIL populations, the simpleM procedure ^49^ was used to estimate the number of independent tests performed. The simpleM approach was modified to deal with missing genotypes introduced by quality control of genotype data. The *fixLength* parameter was tested (from 10 to 300, step size 5) to find the number of independent tests performed (*Meff)*. Using this approach, the number of independent tests was estimated to be 1,546, when the *fixLength* parameter was set to 155. Significance thresholds were calculated using stringent Bonferroni correction to prevent false associations. Significance thresholds were determined as −log_10_(0.01/1,546) = 5.18 LOD to be ‘genome-wide highly significant’ and 4.49 is considered ‘genome-wide significant’ corresponding to genome wide α-level of 0.01 and 0.05, respectively.

QTL confidence intervals were defined when two or more markers show significant association within a 1 Mb region, after which a conservative 2.0 LOD drop from the top marker ^50^ was used; confidence interval start and end positions are defined by the first marker upstream and downstream of the top marker which have a LOD score 2.0 lower than the top marker. An exception to this rule was made when segregation distortion at markers was detected, which was the case for a single novel QTL region (nR3).

### Variance explained

Computing variance explained in an MQM LMM setting is non-trivial, compared to single time point mapping. The biggest nuisance is dealing with the random effects of the model. Our model includes a timepoint|individual random effect, which means that all animals are allowed to have their own start weight at the beginning of the experiment and their own individual growth rate (body weight gain) during the experiment. Including these random effects into the model allowed to better estimate the effect size of the fixed effects of interest. However, the variance absorbed by these random effects counts as unexplained variance in our experimental setup.

Since the random effect needs to be handled as non-explained variance, variance explained is estimated by using the following approach: for each model the estimated fixed effects were used to predict body weights of all animals at each time point. Variance explained was defined as the squared Pearson correlation between the observed body weights and the predicted values.

Since each marker is included in two components, as a main effect (M_(x)_), as well as an interaction term ((M_(x)_:T)), variance explained was computed for each top marker by comparing the variance explained by the model including the main effect and the interaction term, with the variance explained by the model in which both of these components were not included.

### Code availability

LMM-MQM time series mapping code is available at: https://github.com/DannyArends/LMM-MQM-TS. All code is provided under the GNU General Public License version 3.

### Candidate gene analysis

Positional candidate genes were determined for each QTL confidence interval. These gene lists were further prioritized based on their occurrence in several KEGG pathways which encompass the BFMI obesity phenotype on a molecular level. Given that BFMI mice are prone to obesity, KEGG pathways that encompass high fatness and several other features of the metabolic syndrome on a molecular level were selected. KEGG pathways considered can be found in supplemental file 3.

Additionally, a literature study investigating known association of genes within the QTL confidence intervals was performed. Genes were prioritized as candidates if literature evidence was found, e.g. previous association between the gene and body weight, obesity, or type II diabetes.

Further prioritization of candidate genes was performed by the analysis of DNA variations (3’ and 5’-UTR and non-synonymous SNPs) detected within the candidate genes. This was done using DNA sequencing data of the BFMI860 parental strain which was compared to the B6N reference mouse strain available from Ensembl (version GRCm38.6). Based on these mutations and their impact on the gene, further prioritization was performed.

### DNA Sequencing

Paired-end DNA sequencing was performed on the parental line BFMI860 using 150 bp paired end chemistry. 1 µg of genomic DNA was prepared for sequencing using the Illumina PCR free TruSeq protocol. Obtained FastQ data was mapped against the *Mus musculus* mm10 genome using bwa-mem (v.0.7.13) ^51^. Duplicate reads were marked using picard-tools (v.2.4.1) ^52^. Read alignment was checked and prepared for further processing with GATK tools ^53^ using samtools (v.1.3.1) ^54^.

Indel realignment and base quality score recalibration was performed using the GATK (v3.6) following GATK’s best practices workflow ^55^. Afterwards variant calling was performed per sample applying GATK’s HaplotypeCaller in ERC mode yielding variants stored in the variant call file format (vcf) ^56^.

For selected candidate genes known variants were annotated using the dbSNP (version snp138), after which consequences of genomic variants were annotated using the SnpEff tool (version 4.1k) ^57^ available at Ensembl (database version GRCm38).

## Results

LMM-MQM time series mapping detected 5 novel QTL regions. These new regions are designated throughout the results section of this paper as nR*X*, which is short hand for “novel region *X*”, where *X* is the region number as denoted in Table 2.

**Table 2:** Overview of QTLs detected with the LMM-MQM approach. For each region start and stop positions are given based on the conservative 2.0 LOD drop. Effect sizes listed as grams difference at the start of the experiment (columns H and BFMI) and in milligrams/day for the locus x time interaction (columns H/Day and BFMI/Day). All effects are relative to the group of individuals homozygous B6N for the top marker locus (B6N = X and B6N/Day = Y). We list the effect size of the jObes1 locus in the table to allow the comparison of effect size between different loci. The jObes1 locus effects are obtained from the null-model and show a clear dominance pattern in the body weight gain for individuals homozygous for BFMI locus (B6N/Day = Y, H/Day ≈ Y-11 mg/day, BFMI/Day = Y+201 mg/day). We also observe that the estimated intercept is negative (in a dominant way) for the jObes1 locus with homozygous BFMI individuals weighing around 1.44 grams less at the start of weighing (d21) which is consistent with previous findings. All other effect sizes mentioned in the table are from LMM-MQM mapping which include the jObes1 main effect and its interaction with time. The number of individuals in each genotype group is color coded, light green colors represent lower sample size, while dark green represents higher number of individuals.

| Name | Chr | Start | Top marker | Stop | LOD | Number of alleles |  |  | Effect relative to homozygous B6N |  |  |  |  |
| --- | --- | --- | --- | --- | --- | --- | --- | --- | --- | --- | --- | --- | --- |
|  |  |  |  |  |  | B6N | H | BFMI | H | BFMI |  | H/Day | BFMI/Day |
| nR1 | 1 | 149,553,681 | UNC1938399 | 154,868,088 | 7.14 | 83 | 145 | 116 | 0.80 g | 0.41 g |  | -78 mg | -67 mg |
| nR2 | 3 | 26,989,539 | UNC030576333 | 35,953,921 | 5.99 | 46 | 173 | 125 | 0.10 g | 0.42 g |  | 39 mg | -25 mg |
| nR3 | 3 | 49,901,885 | JAX00522656 | 52,973,026 | 5.03 | 60 | 158 | 126 | 0.21 g | -0.08 g |  | 46 mg | 81 mg |
| nR4 | 9 | 86,816,288 | UNC090485124 | 99,363,348 | 6.34 | 171 | 133 | 40 | -0.05 g | 0.49 g |  | -27 mg | -105 mg |
| nR5 | 19 | 37,825,545 | UNC30294194 | 40,410,259 | 4.83 | 81 | 179 | 81 | 0.11 g | -0.37 g |  | 12 mg | 69 mg |
| jObes1 | 3 | 36,481,201 | UNC5048297 | 36,854,743 | 43.23 | 39 | 165 | 140 | -0.07 g | -1.44 g |  | -11 mg | 201 mg |

### Per time point QTL mapping

LM, LMM, and LMM-MQM mapping at individual time points did not identify any new QTL compared to those previously published ^23^ (Figure 1). However, some regions are consistently showing non-significant associations at several time points during the analysis. For example, during the LMM mapping nR5 is consistently detected from day 42 to 70 with a non-significant LOD score between 2 and 4 (Figure 1 - Top). When using LMM-MQM mapping nR1 (day 63 and 70), nR2 (day 49 to 70), and nR4 (day 21, 28, 35, 63, and 70) were detected with non-significant LOD scores between 2 and 4 (Figure 1 - Bottom).

**Figure 1:**
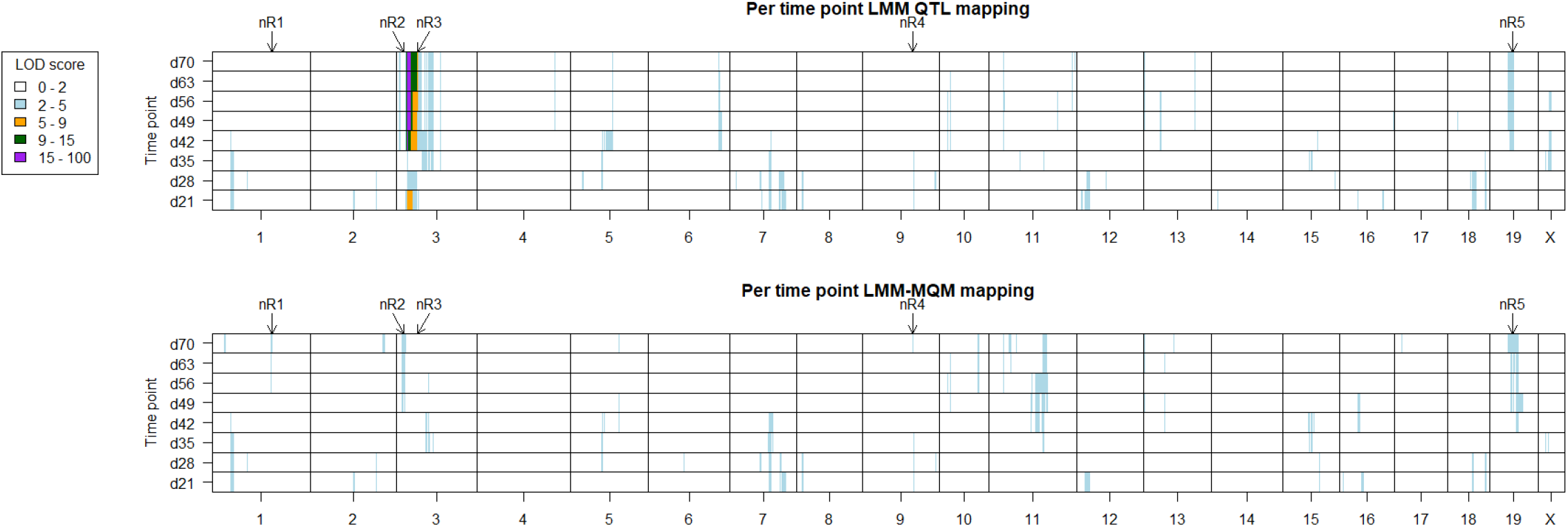
LMM QTL mapping and LMM-MQM mapping for each time point. ***(Top)*** This figure shows the LMM mapping per time point. The *jObes1* locus on chromosome 3 is clearly observed at day 21 (orange) and then from day 42 to day 70 (orange - purple). We can clearly observe the nR5 locus on chromosome 19 from day 42 to 70 with a non-significant LOD score between 2 and 4. Regions nR2 and nR3 are obscured by the strong *jObes1* effect locus. ***(Bottom)*** LMM-MQM mapping per time point, compensated for the strong *jObes1* locus. The intense colors at *jObes1* disappeared, since the effect is absorbed by the *jObes1* top marker covariate in the model. However, no additional significant QTL was detected because statistical power at each individual time point is limited. nR1 is observed on chromosome 1 at day 70, nR2 from day 49 to 70 on chromosome 3, nR3 is not detected in any of the per-timepoint models, and nR4 at days 21, 28, 35, 63 and 70 on chromosome 9, nR5 is consistently detected from day 42 to 70. However, all of these regions did not reach genome wide significance when using separated per time point mapping instead of time series mapping, since LOD scores are in the range of 2 to 4 LOD.

### LMM-MQM time series mapping

An improvement of QTL identification could be obtained by using LMM-MQM time series mapping across all time points. This approach identified five new genome-wide significant QTLs (nR1 to nR5 Table 2), previously undetected during LMM or MQM mapping at individual time points. Variance explained by these novel loci was calculated per time point (Table 3). Between 2.0 % and 3.9 % of the body weight variance can be explained by individual loci. Furthermore, the variance explained by the *jObes1* locus at the end of the experiment (day 70) increased from 25.6 % (single marker mapping) to 36.1 % when using LMM-MQM time series analysis.

**Table 3:** Variance explained by novel QTLs (nR1 to nR5) and *Jobes1* on body weight across all measured time points.

| Age<br>(days) | nR1 | nR2 | nR3 | nR4 | nR5 |  | *jObes1* |
| --- | --- | --- | --- | --- | --- | --- | --- |
| 21 | 1.47 | 0.23 | 1.04 | 0.49 | 0.05 |  | 4.77 |
| 28 | 0.87 | 0.16 | 0.66 | 0.01 | -0.15 |  | -1.35 |
| 35 | 1.70 | 1.09 | 1.65 | 0.82 | 0.46 |  | -0.54 |
| 42 | 2.69 | 0.90 | 2.66 | 2.35 | 1.19 |  | 16.53 |
| 49 | 2.27 | 1.94 | 2.48 | 2.43 | 1.62 |  | 28.83 |
| 56 | 2.82 | 2.76 | 3.01 | 3.85 | 1.40 |  | 34.18 |
| 63 | 2.74 | 1.90 | 2.79 | 3.33 | 1.12 |  | 34.54 |
| 70 | 3.41 | 2.62 | 3.47 | 3.89 | 1.95 |  | 36.05 |

LMM-MQM time series analysis detected two additional QTLs flanking the *jObes1* locus on chromosome 3 (Figure 2). Located proximal of the *jObes1* locus is nR2 (Chr3:26,989,539 - 35,953,921). The nR2 top marker shows that individuals homozygous for the BFMI allele show an increase in growth rate of 23 mg/day compared to homozygous B6N individuals. Heterozygous animals on this locus show a decreased growth rate compared to B6N individuals of −27 mg/day. While these effects look small when expressed on a daily basis this amounts to a difference of ∼2.5 grams between heterozygous and animals homozygous BFMI at the nR2 locus when considering the whole experimental period. The second QTL near the *jObes1* locus (nR3, Chr3:49,901,885 - 52,973,026) located distal of the *jObes1* locus leads to an increase in growth rate for heterozygous individuals (72 mg/day) and homozygous BFMI individuals (43 mg/day) compared to homozygous B6N individuals.

**Figure 2:**
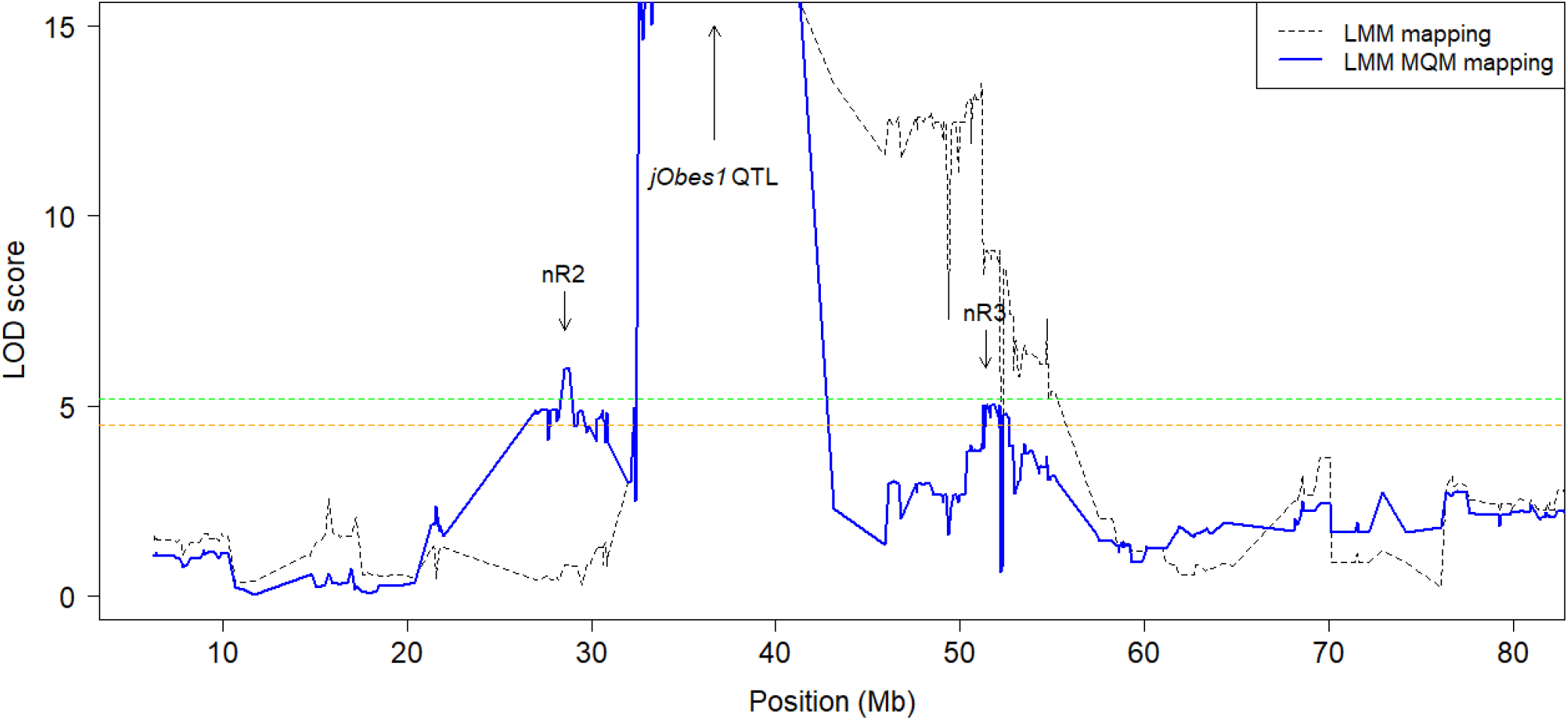
LMM-MQM mapping, detailed view of chromosome 3. After correction for the *jObes1* top marker two new QTLs on chromosome 3 are observed (nR2 and nR3). nR2 shows a genome-wide significant LOD score of 5.99 at the top marker UNC030576333 (Chr3:28,575,896). This locus was not found when mapping without the *jObes1* top marker as a cofactor due to linkage between nR2 and *jObes1*. nR3 shows a genome-wide significant LOD score of 5.03 at the top marker JAX00522656 (Chr3:51,374,784). This locus was also significantly associated before correction for the *jObes1* top marker, but, due to linkage this QTL was hidden by the large *jObes1* QTL. After correction for the *jObes1* QTL, enough evidence remains to detect nR3. Furthermore, a significant drop in LOD scores (Chr3:52,218,893 - 52,370,324) is observable within the confidence interval of nR3 (Chr3:49,901,885 - 52,973,026), this drop is examined more closely in Figure 3.

Interestingly, within the nR3 confidence interval (Chr3:49,901,885 - 52,973,026) a very small region (Chr3:52,218,893 - 52,370,324) was observed that displays a sharp drop in LOD scores (Figure 2 and 3). This small region consisting of 5 markers shows LOD scores below 1.0 LOD, while the markers flanking both sides of this small regions show significant LOD scores above 5.50. Further investigation of this sharp drop in LOD score revealed a segregation distortion within this region. Within this small region, the number of animals homozygous for the B6N allele was severely reduced to below our threshold of 30 individuals per genotype group (Figure 3). The loss of homozygous B6N individuals caused the loss of association in this region, since no significant difference exists between homozygous BFMI and heterozygous BFMI/B6N individuals. Obviously, selection against homozygosity of the B6N allele at this region has taken place. Therefore, we expect that genes located in this LOD-drop region could be responsible for the observed QTL effect.

**Figure 3:**
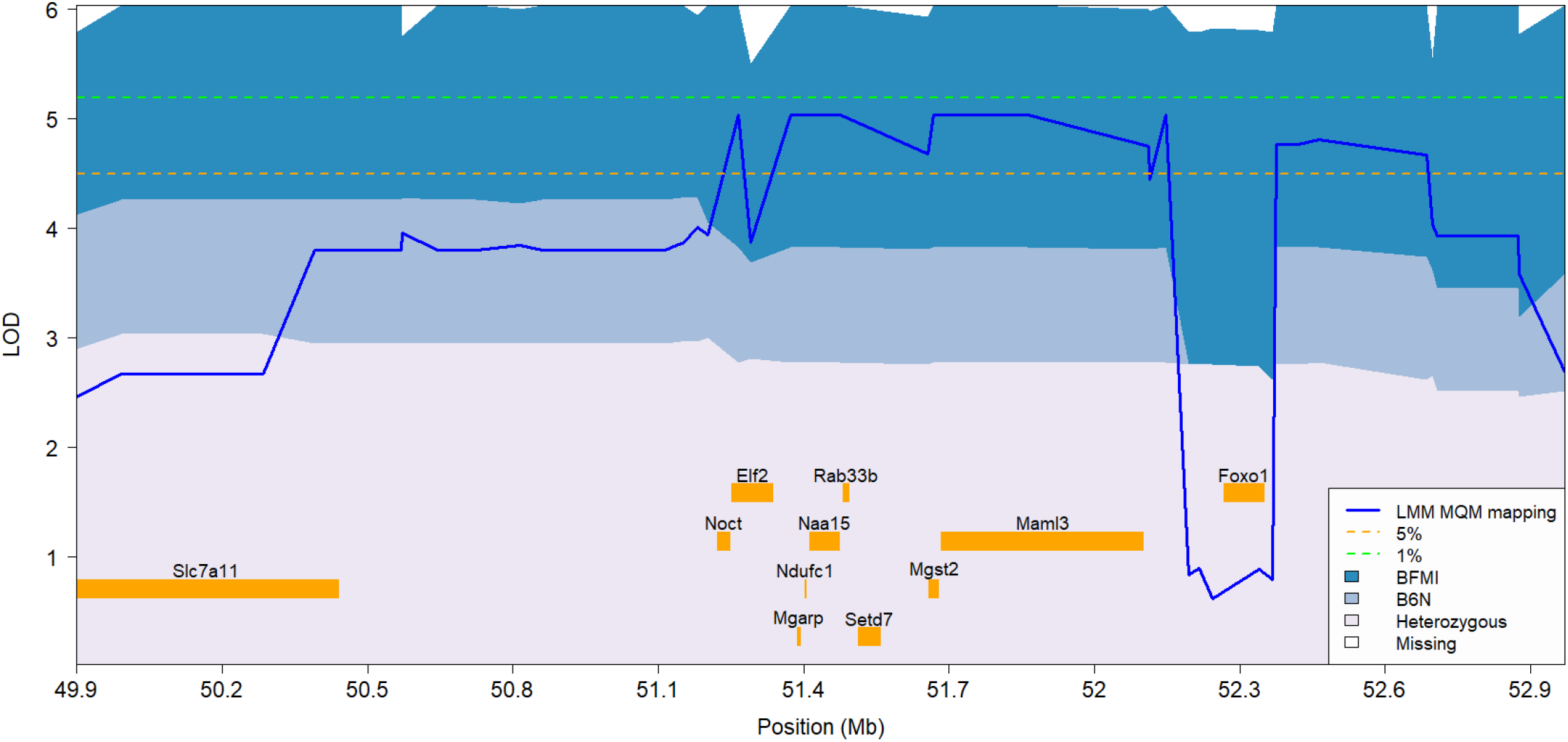
Detail of the nR3 confidence interval showing a sharp drop in LOD scores around *Foxo1*. Distortion of the allele distribution (background coloring of the figure) is observed for 5 SNP markers between 52,218,893 and 52,370,324 base pairs. In the distorted region, a strong increase in the number of individuals homozygous for the BFMI allele is observed (Dark blue, 50% of all individuals in the distorted region), while the percentage of heterozygous individuals stays constant at around 50 %. This leads to a significant distortion from the expected 25:50:25 ratio, which is consistently observed in the surrounding region. The distortion leads to fewer than 30 individuals homozygous for the B6N genotype, causing the B6N/B6N group to be ignored when performing LMM-MQM time series mapping. LOD scores (blue line) sharply drop since the difference between homozygous BFMI and heterozygous individuals is not significant. Located within this drop is only *Foxo1*, a well-known and important regulator within the insulin signaling pathway in the liver.

Surprisingly, this five-marker-region which showed a sharp drop in LOD scores, harbors only one protein coding gene, *Foxo1* (Chr3:52,268,336 - 52,353,221), which has one SNP in the 5’ and six SNPs in the 3’ region of the gene in BFMI mice. The resulting protein (FOXO1) is an important transcription factor which plays a central role in the regulation of glucose metabolism by insulin signaling in the liver ^25^. FOXO1 is also known to be involved in the commitment of a preadipocyte to develop into mature adipocytes ^26^. Furthermore, it was shown that FOXO1 activity decreases lipid droplet formation ^27^ in developing preadipocytes.

### Candidate genes prioritization by KEGG pathways

Most candidate genes were found in the pathways related to insulin signaling (mmu04910) and insulin resistance (mmu04931). KEGG pathway mmu04910 - *insulin signaling* identified five possible candidate genes underlying different nR confidence intervals. Located in the nR2 confidence interval are *Prkci* (Chr3:30,995,747 - 31,052,959) and *Pik3ca* (Chr3:32,397,671 - 32,468,486) ^28^. The previously discussed *Foxo1* (Chr3:52,268,336 - 52,353,221) gene is located in the nR3 confidence interval. *Pik3cb* (Chr9:99,036,654 - 99,140,621) is located in nR4, and *Sorbs1* (Chr19:40,294,753 - 40,513,779) in nR5. KEGG pathway mmu04930 - *Type II diabetes mellitus* identified another possible candidate gene: *Cacna1e* (Chr1:154,390,731 - 154,884,501) located in nR1. In addition, KEGG mmu04950 - *Maturity onset diabetes of the young*, revealed another candidate gene: *Slc2a2* (Chr3:28,697,903 - 28,731,359) in nR2. mmu04923 - *Regulation of lipolysis in adipocytes* adds *Ptgs2* (Chr1:150,100,031 - 150,108,227) located in nR1 as another candidate gene. In nR4 KEGG mmu04973 - *Carbohydrate digestion and absorption* identified *Atp1b3* (Chr9:96,332,655 - 96,364,442) as a further candidate.

### Candidate genes per region

Within the nR1 region (Chr1:149,553,681 - 154,868,088), a total of 34 protein coding genes are located. The KEGG pathway search identified one possible candidate (*Cacna1e*) while literature search found another three candidates previously implicated in the regulation of body weight: *Pla2g4a* associated with resistance to high fat diet induced obesity and insulin sensitivity ^29^, *Ptgs2* which was shown to be epigenetically dysregulated in diabetes-prone bicongenic B6.NODC11b x C1tb mice ^30^, and *Prg4* which upon weight loss in human showed a significant long-term decrease in blood plasma ^31^. Furthermore, *Prg4* deficiency was found to protect against glucose intolerance and fatty liver disease in diet-induced obese mice ^32^.

Out of the 45 protein coding genes that are located within the nR2 region (Chr3:26,989,539 - 35,953,921) two genes (*Prkci and Pik3ca*) are in the KEGG *insulin signaling pathway*. The gene *Slc2a2*, which is part of two KEGG pathways (mmu04931 - *MODY*, and mmu04973 - *Carbohydrate digestion and absorption*), is also located within the nR2 QTL region. Literature yielded another four candidates: *Cldn11*, one of the five most regulated transcripts in the pancreatic islets of the obesity sensitive New Zealand Obese (NZO) strain compared to the B6N-ob/ob mice protected from diabetes due to increased β-cell production ^33,34^. Furthermore, *Kcnmb2* and *Kcnmb3*, shown to have genetic variants which are associated with insulin resistance, and are modified by dietary (polyunsaturated) fat content ^35^ can be regarded as candidates. The last candidate from literature is *Sox2*, often discussed as possible candidate gene in type II diabetes, but its role and association are still unclear.

The nR3 region (Chr3:49,901,885 - 52,973,026) contains only 11 protein coding genes. Located in this region is the *Foxo1* gene which was prioritized due to literature and its prominent role in the KEGG *insulin signaling pathway*. Furthermore, *Foxo1* contains one SNP in the 5’ and six SNPs in the 3’ region of the gene in BFMI mice. The second candidate for this region *Setd7* comes from literature. *Setd7* was found to be significantly downregulated after caloric restriction in rhesus macaques ^36^, and has been implicated to be involved in the epigenetic regulation of macrophage activation in type II diabetes ^37^. The third candidate gene in this region is *Rab33b* one of the interaction partners of TBC1D1/4 which is involved in insulin-dependent GLUT4 trafficking ^38,39^.

A total of 66 protein coding genes are located within the nR4 region (Chr9:86,816,288 - 99,363,348). Besides the *Pik3cb* gene (the interaction partner of *Pik3ca*, located within nR2) which was prioritized due its KEGG pathway, two additional candidate genes were detected within the nR4 region: *Atp1b3* which was annotated to the insulin secretion pathway in gene ontology, and *Mrap2* which has been associated with severe obesity in mammals ^40^.

The nR5 region (Chr19:37,825,545 - 40,410,259) contains 28 protein coding genes, three of which can be considered as possible candidate genes: *Ffar4*, involved in brown fat thermogenesis ^41^, *Rbp4*, an adipokine that contributes to insulin resistance in the AG4KO mouse model ^42^, and *Sorbs1* in which the T228A polymorphism in humans has been associated with obesity and type II diabetes ^43^.

### DNA Sequencing

In total 25 candidate genes within the five new QTL confidence intervals were prioritized for functional relevance to the BFMI phenotype. DNA sequence variants for these genes occurring in the BFMI860 founder line compared to B6N are summarized in supplemental file 4. Here only mutations with profound effects are mentioned for brevity.

Most striking is the 50 amino acid deletion found in the *Prg4* gene (nR1) in the BFMI860 founder line. This large deletion of 50 amino acids out of 1221, removes amino acids 600 to 650 in exon 7, and most likely results in a non-functional protein. This large deletion combined with the previously discussed literature evidence makes *Prg4* the top candidate for the nR1 region.

*Atp1b3* located in nR4 has four non-synonymous SNPs (S127T, V247F, H250R, and R277Q) in the BFMI860 founder line. These mutations are found to be heterozygous in the BFMI860 line despite inbreeding, while SNPs outside of this gene are found to be homozygous. As such, our hypothesis is that these three mutations lead to a non-functional protein, which when both alleles are affected cause (embryonic) lethality. The occurrence of these three non-synonymous SNPs elevates *Atp1b3* to the top candidate for the nR4 region.

Three non-synonymous mutations (S200P, T236A, and P309L) were detected in the *Sorbs1* gene (nR5). Since, the mutations are homozygous in the BFMI860 founder strain, the resulting protein is likely functional. However, the non-synonymous SNPs might cause protein kinetics to be different between the BFMI and the wildtype B6N protein variant and thereby contribute to the observed phenotypic differences between the strains.

## Discussion

When dealing with interacting and/or correlated effects, it is often more statistically powerful to treat them as one categorical variable. Time series modeling is complicated and needs to be performed carefully in a stepwise fashion, adding effects stepwise to the model, while comparing models and reasoning about them at every step of the modeling process. An example of this is litter size and litter number which both are significant factors in the examined AIL population, but also have a weak but significant interaction (p = 0.098). In such a situation, it is better to model these two linear covariates as a single combined categorical variable “litter type” (which is the combination of litter number and litter size). This significantly improved the resulting model, and removes the need to include an interaction term, which would add to model complexity. Furthermore, including the categorical variable “litter type” increases the available number of degrees of freedom remaining for marker - phenotype association analysis.

The resulting LMM-MQM model contains many factors which may lead to spurious associations due to small sample size in one of the genotype groups. Therefore, we applied very stringent quality control to genetic markers. This leads to a reduction in markers that can reliably be used in mapping time series data. For example in this study 17,971 markers were found to be reliable for usage during LMM-MQM time series mapping, versus 22,164 markers that were used during our previous research ^23^ where single marker, single timepoint QTL mapping was performed. This reduction in the number of suitable markers decreases the chance of false associations while simultaneously increasing reliability for newly detected loci.

Time series data on body weights are often collected during experiments, however, when QTL mapping this time series structure of the data is often not fully exploited. QTLs are mapped at individual time points, resulting in less power to detect small effect QTLs or QTLs acting continuously in a time dependent manner. The combination of LMM and MQM mapping methods allows the more fine-grained detection of QTL by modeling all time points together and compensating for significant genetic effects. This led in our study to the identification of five additional loci affecting body weight in young mice.

Within each identified region known and unknown candidate genes can be responsible for the phenotypic alterations. The identification of regions without known candidates are interesting as they allow discovery of novel candidate genes contributing to the development of obesity over time. Finding several prior known associations, on the other hand, demonstrates that LMM-MQM mapping improves the association of genetic factors with body weight, since standard LM and LMM mapping failed to detect these associations.

Currently more and more phenotypic data is collected across a wider time span ^44^. This wealth of temporal data combined with the decrease of costs for genotyping and sequencing allows for more sensitive modeling of genetic effects underlying these phenotypes in a wide range of species. However, this also requires the development of novel methods and algorithms to analyze this high dimensional data and exploit the temporal structure of this data ^5^.

## Conclusions

QTL mapping using the MQM approach accounts for known genetic effects and can be combined with LMM time series mapping for more statistical power. It allows to model repeated measurements of a trait on multiple individuals by modeling the full time series, in contrast with traditional single time point methods that treat time points individually. Using a full model with all available data, five additional significant genomic regions contributing to body weight variance were detected. While the LMM approach individually allows detection of additional loci on different chromosomes, complementing the LMM approach by MQM the new method was able to compensate for the large effect of the *jObes1* which led to the detection of additional QTLs, but also to the dissection of flanking regions of the *jObes1* locus.

LMM-MQM timeseries mapping is a generic approach which models complex high dimensional data, compensates for known genetic effects, and takes advantage of the temporal structure of phenotypic data.

## Declarations

### Availability of data and materials

All data generated during this study are included in this published article and its supplementary information files. R code (as well as tab separated input data) to perform the LMM-MQM time series mapping are available at: https://github.com/DannyArends/LMM-MQM-TS. All code is provided under the GNU General Public License version 3. DNA sequencing data were deposited at the NCBI Sequence Read Archive (SRA) under BioProject ID: PRJNA717237.

### Competing interests

The authors declare that they have no competing interests.

### Funding

The project was funded by the DFG (BR 1285/12).

### Author contributions

DA - Method design, data analysis, and wrote the manuscript

DH - Candidate literature gene analysis, method discussion, and manuscript revisions

SK - Data collection, and method discussion

SH - Animal handling, and data collection

SL - Method design (variance explained), data analysis, and manuscript feedback

PK - Method design (variance explained), designed figure 3, and manuscript feedback

MD – Tested the method in another population, helped generalize the method

MM - Method discussion, and manuscript feedback

PP - Method discussion, and manuscript feedback

GB - Experimental design, supervision, method discussion, and manuscript feedback and revisions

## Acknowledgments

Not applicable.

## Supplemental files

Supplemental file 1 - Phenotypes and genotypes of the AIL individuals

Supplemental file 2 - Model selection between litter number, size, and type

Supplemental file 3 - KEGG pathways considered associated with the BFMI obesity phenotype

Supplemental file 4 - Genes in novel QTL confidence intervals, candidate gene prioritization and mutations detected by DNA sequencing of the BFMI860 founder line

